# Identity-by-descent mapping using multi-individual IBD with genome-wide multiple testing adjustment

**DOI:** 10.1101/2025.01.28.635369

**Authors:** Ruoyi Cai, Sharon R. Browning

## Abstract

We present an identity-by-descent mapping approach to test the association between genome-wide loci and complex traits. Our method evaluates whether levels of genetic similarities at specific genomic locations, captured by local relatedness matrices derived from multi-individual IBD sharing, are associated with phenotypic variation in complex traits. In addition, we propose an approach to adjust for multiple testing in genome-wide IBD mapping scans based on the correlation structure between test statistics across the genome. Through simulation studies, we demonstrate that our test has a well-controlled genome-wide type-I error rate and superior power to detect rare and untyped variants compared to standard single-variant tests. We applied our method to systolic blood pressure data from White British individuals in the UK Biobank.

## Introduction

With the development of sequencing technology, there has been a surge in understanding the association between genetic variants and complex traits over the past two decades. Historically, there are two major classes of methods for association analysis: linkage mapping and genome-wide association studies. Linkage mapping identifies regions of the genome that are coinherited with a trait within families, which is particularly powerful for detecting rare variants.^1-5^ However, linkage mapping is often limited by its low resolution and requirement for family data. In contrast, as a more commonly used method at present, genome-wide association studies (GWAS) can analyze common genetic variants across large, unrelated populations, offering high resolution and the ability to detect associations with complex traits at a genome-wide level.^6-8^

While single-variant tests in GWAS are powerful for detecting common variants, they often struggle to detect structural variants or rare variants that are more likely to be population-specific, untyped in the genotype data, and have small individual effect sizes.^9-11^ Variant-set tests improve upon single-variant tests by aggregating effects across multiple variants within a genomic region or a set of related genes, thus increasing power to detect rare variants.^12-15^ However, these methods often rely on pre-defined models about the genetic architecture underlying complex traits, the sparsity of causal variants, and the choice of weights on the effects of variants. In comparison, population-based identity-by-descent (IBD) mapping offers a complementary approach for linkage mapping, GWAS, and variants-set tests, providing a more flexible and comprehensive framework for association testing.

IBD mapping identifies genomic regions containing potential causal variants by searching for genomic positions where levels of IBD sharing are associated with phenotypic variation in a group of individuals for a trait of interest.^16-26^ By leveraging long stretches of IBD segments shared between individuals, IBD mapping can capture the combined effects of co-occurring proximate rare alleles, even when individual variants have small or modest effects.^18,19,27^ In addition, IBD mapping approaches can indirectly recover signals of untyped rare variants, structural variants, or population-specific variants tagged by IBD haplotypes.^16-19,26^ Furthermore, IBD mapping is not reliant on assumptions about the underlying genetic structure or models on the effects of causal variants, which contributes to its applicability and robustness in analysis of complex traits with unknown or intricate genetic architectures. ^12-15^

Previous IBD mapping studies have mostly focused on using IBD segments shared between pairs of individuals.^16,18,21-26^ However, IBD information can extend beyond the pairwise level by considering clusters of haplotypes that are all IBD with each other.^17,20,28,29^ Using IBD haplotypes shared among groups of individuals may enhance power for IBD mapping due to more accurate IBD calls and the additional information provided by specific IBD group effects that are ignored in pairwise IBD analysis. Nevertheless, multi-individual IBD mapping has not been widely used to study variants underlying complex traits due to challenges in identifying IBD clusters in biobank-scale studies and in designing versatile models to incorporate IBD cluster information for association testing.^17,20^

In this work, we develop an IBD mapping test leveraging multi-individual IBD sharing between distantly related individuals in large, outbred populations. Motivated by variance component models in linkage analysis, we construct local relatedness matrices from multi-individual IBD sharing to quantify genetic similarities at consecutive genomic locations. Using a likelihood ratio framework, we test for associations between genomic regions and a complex trait of interest by evaluating whether local genetic similarities significantly contribute to the phenotypic variation of the trait. Additionally, we develop an approach to adjust for multiple testing based on an analytical formula for the correlation between test statistics across the genome. Through simulation studies, we demonstrate that our test maintains a well-controlled family-wise type-I error rate and exhibits superior power to detect the effects of rare and untyped variants in SNP array data compared to standard single-variant tests. Applying this method to White British individuals in the UK Biobank, we identified signals associated with systolic blood pressure in this cohort.

## Methods

### Variance component test

We use a quantitative traits model that is commonly used in linkage analysis.^1-3,5,30^ Denote ***Y*** as the vector of quantitative trait values observed from a population sample. We model the trait values with the following linear mixed-effects model:

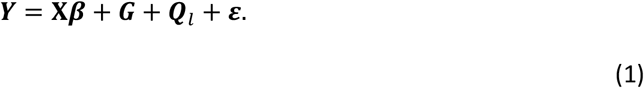

In the above formula, **X** is a matrix of fixed-effects covariates with effect sizes ***β. G*** and ***Q***_***l***_ are random-effects terms that represent the genome-wide additive effect and the location-specific effect at position *l* respectively, and ***ε***is the environmental effect. The genome-wide additive effect, ***G***, is assumed to follow a multivariate normal distribution 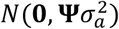, where **ψ** is the genome-wide relatedness matrix that can be estimated from genome-wide IBD sharing. We refer to **ψ** as the global IBD matrix. The location-specific effect ***Q***_*l*_ at position *l* is assumed to follow the multivariate normal distribution 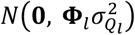, where **Φ**_*l*_ is the location-specific relatedness matrix that characterizes the proportion of alleles shared IBD between each pair of individuals at a specific genomic location *l*. We refer to **Φ**_*l*_ as the local IBD matrix at position *l*. The environmental effect is assumed to be independent between individuals and follow a normal distribution with mean 0 and variance 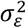.

Given the mixed-effects model in (1), the phenotypic variance of the quantitative trait *Y* is represented by the sum of three variance components:

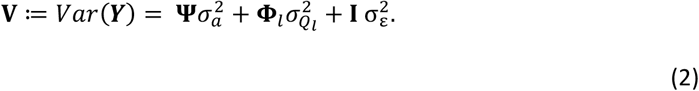

The parameters 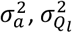 and 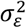 represent the effect sizes of the three variance components. The structuring matrices **ψ, Φ**_*l*_, and **I** predict the covariances among individuals attributable to the effect of each variance component, respectively. Testing whether a locus at position *l* is associated with the quantitative trait is equivalent to testing the variance component hypothesis 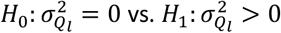.

Many different forms of test statistics have been proposed for variance component tests. In this study, we use the log-of-odds (LOD) score^3,31^ as the test statistic to take advantage of the maximal information considered under the likelihood ratio framework. To test 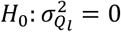 vs.

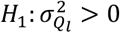, we consider the test statistic *W*_*l*_ defined as:

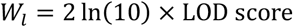

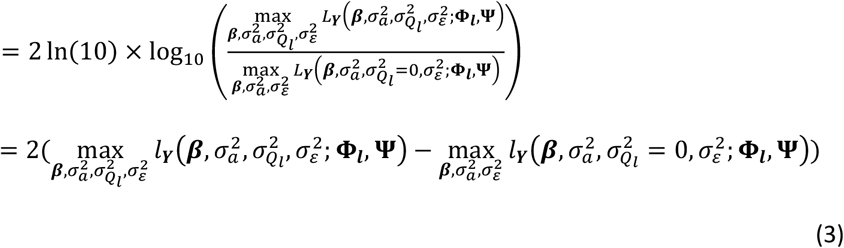

where 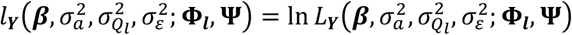 is the restricted model log-likelihood that takes the following form, with the variance matrix **V** defined in (2):

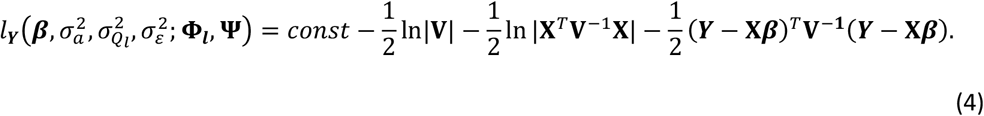

To calculate the test statistic, we find restricted maximum likelihood estimators (REML) of the fixed effect ***β*** and variance effects 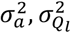 and 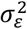. Under the null hypothesis, the distribution of *W*_*l*_ is a 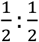 mixture of a point mass at 0 and a χ^2^ distribution with 1 degree of freedom (Appendix A).^3^

The application of the likelihood ratio statistic in variance component tests faces two major challenges. First, without closed-form solutions for REML estimates of variance components, an efficient and accurate numerical optimization algorithm is essential. Second, calculating the log-likelihood for multiple iterations during optimization can become computationally expensive with large sample sizes (*N*), as matrix inversion typically has a complexity of *O*(*N*^3^). To address these issues, we use the BFGS algorithm^32-36^ from the SciPy optimize^37^ library in Python due to its robustness and efficiency in handling large-scale optimization problems. Additionally, we stored the global and local IBD matrices as sparse matrices using the data structures and functions provided by the SciPy sparse library, ensuring efficient storage and manipulation of the variance matrix. Furthermore, to expedite calculation while ensuring numerical stability, we avoid direct matrix inversion by first performing sparse matrix Cholesky decomposition and then conducting forward substitution, with utilities implemented in the scikit-sparse library.^38,39^

### Multi-individual identity-by-descent inference

In this study, we focus on haplotypes that are identical by descent across multiple individuals instead of between pairs of individuals, which provides more detailed information about IBD sharing across the genome. Previous approaches usually identify multi-individual IBD by clustering pairwise IBD haplotypes. As the number of pairwise IBD segments scales quadratically with sample size, such approaches would require an impractically high amount of memory and time on biobank-scale dataset.^17,20,29^

In this study, we use the recently developed software ibd-cluster that scans locations across the genome to directly cluster haplotypes that are IBD at each specific genomic location without identifying pairwise IBD segments.^28^ This algorithm first assigns each haplotype to an individual cluster and then merges clusters when a pair of haplotypes, one from each cluster, share the same allele sequence in a genomic region of at least *L* cM that contains the markers of interest and extends at least *T* cM (*T* < *L*/^2^) in both direction from the markers of interest. We refer to *L* as the haplotype length threshold and *T* as the trimming threshold. Due to IBD transitivity, only clusters for pairs of haplotypes that are adjacent in the positional Burrows-Wheeler Transform sorting need to be merged, so this approach scales linearly with the number of individuals in the dataset in terms of computation time, memory requirements, and output size. This makes it feasible for using with very large datasets, such as those in biobanks.

At a given genomic location, a group identifier representing the IBD state is assigned to each of the two haplotypes for each individual in the sample. Haplotypes sharing the same group identifier are identical by descent with each other at the given location. This format is highly efficient for computing local relatedness matrices, as it avoids iterating through a potentially quadratic number of IBD segments relative to the sample size. Instead, computations are performed directly on a vector of haplotype cluster indices, which scales linearly with the sample size.

Using smaller haplotype length thresholds *L* or smaller trimming thresholds *T* to detect multi-individual IBD will lead to inclusion of shorter shared haplotypes in clustering, which will potentially lead to larger multi-individual IBD clusters. As a result, the local IBD matrix will tend to be less sparse, and also more likely to include false-positive IBD. In contrast, using larger *L* and *T* to detect multi-individual IBD imposes a more stringent standard when clustering IBD haplotypes, and thus the resulting multi-individual clusters will tend to contain longer but fewer segments, leading to a sparser local IBD matrix. Through simulation studies, we evaluate how different choices of *L* and *T* for multi-individual IBD inference affect the performance of our IBD mapping test.

### Global and local IBD matrices

The global IBD matrix **ψ** is a symmetric matrix, with 1’s on the diagonal, and the (*i, j*)-th entry being the coefficient of relatedness between the *i*-th and the *j*-th individuals, which equals twice the kinship coefficient of this pair of individuals. To construct the global IBD matrix, we infer pairwise IBD segments using the software hap-ibd^40^ from phased genotype or sequence data, and estimate kinship coefficients using the software IBDkin.^41^ IBDkin calculates kinship coefficients based on the proportion of the genome shared between pairs of individuals with 1 or 2 IBD haplotypes. Unlike averaging local relatedness matrices across genomic locations, which can be computationally intensive and susceptible to noise from sparse or unevenly distributed local IBD data, IBDkin leverages genome-wide information to yield a robust and efficient estimate of relatedness. To maintain the sparsity of **ψ**, we set kinship coefficients below 0.044 (corresponding to relationships beyond third-degree relatives) to zero.

The local IBD matrix **Φ**_*l*_ characterizes the genetic similarities between individuals at a specific position *l* on the genome. We utilize multi-individual IBD inferred using the ibd-cluster software to obtain the localized relatedness coefficient at position *l* between each pair of individuals in the local IBD matrix. We summarize the proportion of each individual’s chromosome in each multi-individual IBD group at the given location *l* with a matrix **A**_*l*_. We define **A**_*l*_ as a *N* × *K*_*l*_ matrix, where *N* is the number of individuals in the sample and *K*_*l*_ is the total number of IBD clusters at the given location *l*. The (*i, k*)-th entry of **A**_*l*_ is the proportion of individual *i*’s chromosomes (0, ½, or 1) that are assigned to IBD group *k* at locus *l*.

The local IBD matrix at genomic location *l* is then obtained as 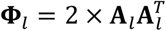. This means that we sum the proportion of IBD haplotypes over all clusters at *l* for each pair of individuals. As a result, **Φ**_*l*_ is a symmetric matrix with diagonal of 1’s and the (*i, j*)-th entry as the proportion of alleles shared IBD by individuals *i* and *j* at position *l*, which would be non-zero only if individuals *i* and *j* have haplotypes in the same IBD cluster. Local IBD matrices constructed in this way are guaranteed to be positive (semi)definite, which ensures that Cholesky decomposition can be performed on the variance matrix in our optimization algorithm to efficiently optimize the model likelihood.

### Genome-wide multiple testing adjustment

To access the significance of test results in IBD mapping scan across the genome, correcting for multiple testing is essential. The commonly used Bonferroni threshold would be conservative for IBD mapping tests due to correlations between tests at nearby locations that are covered by the same IBD segment. To obtain an appropriate genome-wide significance threshold, we need to consider the joint distribution of test statistics when performing IBD mapping at a series of locations across the genome.

Consider standardized test statistics {*Z*_*l*_} that are approximately normally distributed and follow an Ornstein-Uhlenbeck (OU) process under the null hypothesis of no genetic associations. The mean *E*[*Z*_*l*_] = 0, and the covariance between any pair of test statistics 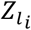 and 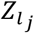 is given by 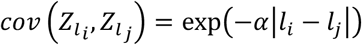.^42-44^ The parameter α > 0 is the rate of correlation decay as genomic distances increases between test statistics, and we refer to it as the decay parameter of the OU process.

We show in Appendix B that the IBD mapping statistics 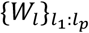 for testing genomic location *s l*_1_, *l*^2^, … , *l*_*p*_ can be define in terms of standard normal {*Z*_*l*_} as:

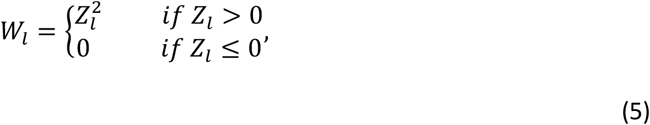

If the 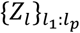 form an OU process, then we refer to 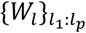 as the modified OU process. We expect that the IBD mapping statistics will be approximately memoryless as a process along the genome, so that the modified OU process may provide a good approximation to the joint distribution of the test statistics. In what follows, we assume that the IBD mapping statistics do indeed follow a modified OU process.

Define *d* = |*l*_*i*_ − *l*_*j*_| and *ρ*(α, *d*) = exp(−α*d*). It can be shown that at any locations *l*_*i*_ and *l*_*j*_ that are spaced at intervals of length *d*, the correlation between 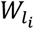 and 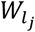 depends only on the decay parameter α and the distance *d* (Appendix B):

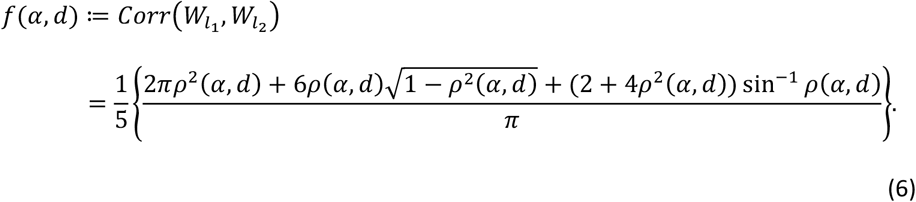

Based on (6), we can estimate α from observed correlations between LOD scores under the null hypothesis. To do so, we simulate phenotype values under the null hypothesis and use pairs of LOD scores obtained at any two locations spaced at distance *d* to calculate the empirical correlation 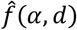, and then estimate 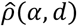 by solving 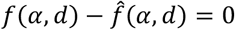 for a given *d*. Since *ρ*(α, *d*) = exp(−α*d*), we have − log(*ρ*(α, *d*)) = α*d*. Hence, α can be estimated by the slope of a linear regression line without intercept between – 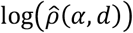 and *d* at a series of different values of *d*.

Given 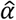, we implemented a Monte Carlo approach find the significance threshold for genome-wide multiple testing based on the distribution for max{*W*_*l*_}. We simulate observations 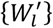 from the modified OU process, starting from *l* = 0 and incrementing *l* by the same space between test locations selected in genome-wide IBD mapping, until *l* reaches the end position on the genome. The simulated process 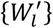 represents a single replicate of the genome-wide IBD mapping test under the null hypothesis. An empirical distribution for 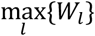 can be obtained by repeating the simulation thousands of times and recording 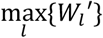 from each replicate. We take the 95% quantile from this empirical distribution as the 95% significance threshold of genome-wide IBD mapping on the given data. The detailed algorithm to simulate the modified OU process along the genome is given in Appendix C. We note that the genome-wide multiple testing correction depends on the specific settings used to detect multi-individual IBD, because the length distribution of IBD haplotypes considered in the algorithm will affect the covariance between test statistics.

### Analysis pipeline

The required input data for our IBD mapping test include phased genotype sequence or array data, a quantitative trait of interest, fixed-effect covariates if applicable, and a genetic map.

As the first step, we construct a global IBD matrix and location-specific relatedness matrices. For the global IBD matrix, we use kinship coefficients that are estimated by the IBDkin program based on pairwise IBD segments identified from the sample using the hap-ibd program, as described above. For the hap-ibd analysis on sequence data, we set the minimum seed length to 0.5 cM, the minimum extension length to 0.2 cM, and the minimum output segments length to 2 cM. We also exclude rare variants by setting the minimum minor allele count filter to 100. For the hap-ibd analysis on SNP array data, due to the lower marker density, we set the minimum seed length to 1 cM, the minimum extension length to 0.1 cM, the minimum number of markers in a seed IBS segment to 50, and the minimum output segments length to 3 cM. Other parameters are left at their default values. When estimating kinship coefficients, we use all default parameters of the IBDkin program.

Next, we run the ibd-cluster software to obtain multi-individual IBD clusters. In our simulation studies, we applied different haplotype length thresholds *L* at 2 cM or 3 cM and different trimming thresholds *T* at 0.25 cM, 0.5 cM, or 0.1 cM to investigate how different IBD clustering thresholds affect the performance of our test. When analyzing UK Biobank data, we set *L* = ^2^ cM and *T* = 0.5 cM, which is the setting that we found through our simulation studies to give good power while produce reasonably sparse local IBD matrices that are computationally feasible for large-scale data.

We then select a collection of locations across the genome as candidates to compute the local IBD matrices and perform the IBD mapping test. The test locations do not need to be directly available in the array data or in the IBD cluster output, as we can simply use the nearest location to the testing position in the IBD cluster output to perform the test. As IBD states do not change immediately across nearby markers, we recommend running tests at 0.1 cM intervals when considering IBD clusters based on haplotypes that are at least 2 cM long.

For the simulation studies, and to estimate the genome-wide multiple testing threshold in the UK Biobank data, we ran IBD mapping tests at 0.1 cM intervals. When analyzing phenotypes from the UK Biobank, to save computational resources given the large sample size, we chose a two-step approach. On each chromosome, we first ran IBD mapping test at locations spaced 1 cM apart across the genome, and we identified the top 10 locations ranked by ascending test p-values. We then zoomed into 1 cM regions centered around these locations by conducting finer-scale tests at 0.1 cM intervals.

Our method is also applicable to genomic regions in addition to single genomic locations. We compute the local IBD matrix over a genomic region by averaging the local IBD matrices computed at a few selected locations within the given region. In the simulation studies for power estimation, we tested each 0.05 cM region containing simulated causal variants using the local IBD matrix computed by averaging the IBD matrices at the start and the end positions of the region.

Finally, we compare test p-values to the genome-wide significance threshold. To derive the multiple testing adjustment for genome-wide IBD mapping, we first estimate the decay parameter α of the modified OU process formed by IBD mapping statistics across the genome under the null hypothesis, as described above. Null phenotypes are simulated as the sum of independent random variables drawn from the standard normal distribution and random variables drawn from the multivariate normal distribution with mean 0 and covariance matrix 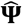, where 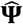 is the global IBD matrix estimated from the data. We conduct IBD mapping tests at 0.1 cM intervals across the genome and calculate the empirical correlation 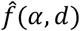 between pairs of test statistics spaced at *d* cM for a sequence of *d* from 0.1 cM to 1 cM, spacing at 0.1 cM intervals. We then estimated 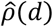 from 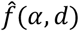 based on equation (6) for each *d*. Finally, we fit a regression on –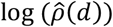 against *d* and use the slope of the fitted line as the estimate 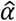 for the decay parameter α.

We perform a bootstrap analysis across chromosomes to obtain confidence intervals for 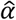. Specifically, we set the number of bootstrap replicates to 10,000, and we resample chromosomes with replacement in each replicate. We take all pairs of test statistics spaced at *d* cM within each chromosome, and combine pairs over all sampled chromosome to calculate the sample correlations and estimate the decay parameter 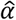. We calculate the 2.5^th^ and 97.5^th^ percentile of 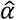 across all bootstrap replicates to construct the 95% confidence interval for the estimated decay parameter 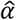.

We then use 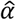 as the decay parameter to simulate observations at 0.1 cM intervals from a modified OU process that spans the same distance as the total size of the genome for 10,000 replicates, and we take the 95% quantile of maximum test statistics from all simulations as the genome-wide significance threshold.

### Studies on simulated data

We used msprime^45,46^ to simulate whole-genome sequence data for 5,000 individuals, with each genome consisting of 30 chromosomes, all 100 cM in length. The demographic model for the simulation, adopted from previous studies, resembles the demographic history of the UK population.^47^ In the msprime simulations, mutation occurred at a rate of 10^−8^ per base pair per generation, and recombination at a rate of 10^−8^ per base pair per meiosis.

We estimated the genome-wide type I error rate in the simulated sequence data by simulating 1000 replicates of phenotypes without genetic associations on random samples of 1000 individuals from the original dataset. On the simulated sequence data, we generated phenotypes following ***Y*** = ***g*** + ***ε***, where ***g*** follows a multivariate 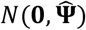 distribution, with 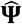 being the genome-wide relatedness matrix constructed from estimated kinship coefficients of all pairs of individuals in the sample, and *ε* is a vector of standard normal observations. For each replicate, we performed genome-wide IBD mapping at 0.1 cM intervals and obtained a genome-wide significance threshold based on the observed test scores. We calculated the genome-wide type I error rate as the proportion of replicates that contain at least one test score achieving the corresponding genome-wide significance threshold.

We estimate the power of our test by simulating phenotypes associated with four types of causal variants using simulated sequence data. Variants in the sequence data are classified based on minor allele frequencies (MAF) as common (MAF > 10%), low-frequency (1% < MAF < 10%), rare (0.05% < MAF < 1%), and ultra-rare (MAF < 0.05% but appearing at least once in the data). We simulate 1000 replicates of phenotypes associated with a 0.05 cM region containing *k* causal variants in each of these four classes.

For phenotypes associated with common or low-frequency variants, we consider regions that contain at least 4 variants of the target class, and designate half of them as causal. For rare and ultra-rare variants, we consider regions that contain at least 8 variants of the target class, and randomly select a quarter of them as causal. The simulated phenotype for the *i*^*th*^ individual is constructed as 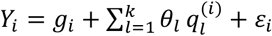, where *g*_*i*_ and *ε*_*i*_ are the *i*^*th*^ observations of the vectors ***g*** and ***ε***that are defined as above, 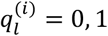 or 2 is the number of copies of the minor allele at marker *l*, and the effect size of each causal marker depends on its minor allele frequency as 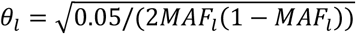.

From the simulated sequence data, we generate a simulated SNP array dataset for each chromosome by excluding all variants with MAF below 1% and all variants associated with the simulated phenotypes, and then randomly selecting 30,000 variants per chromosome. We evaluate the power of our test to detect variants associated with the simulated phenotypes using both the full simulated sequence data, where all genetic variants are directly available, and the simulated SNP array data, where none of the causal variants are directly present.

The power of our IBD mapping test is calculated using both the Bonferroni threshold (0.05/^3^0000 = 1.67 × 10^−6^) and the genome-wide significance threshold derived from our proposed multiple testing adjustment. On simulated sequence data, we also conducted the SKAT^13^ test using default parameter settings to benchmark the performance of our IBD mapping test. On the SNP array data, we compare the performance of our test to the traditional single-variant test used in GWAS. Within each 0.05 cM region containing causal variants, we test every marker in the SNP data using simple linear regression between the phenotype value and the copy number of minor alleles. The minimum p-value across all tests within a region is used as the overall single-variant test p-value for that region. For the SKAT test, power is calculated using the significance threshold 10^−6^, as recommended in Wu et al. (2011) for genome-wide testing.^13^ For the single-variant test, power is calculated using the traditional GWAS significance threshold 5 × 10^−8^, which is approximately 0.05 divided by the total number of markers in our simulated SNP array data.

### UK Biobank data analysis

The UK Biobank SNP-array data were collected on half a million UK participants using the UK Biobank Axiom array that assays approximately 850,000 genetic variants across genome.^48^ We analyzed the systolic blood pressure of White British UK Biobank individuals.

We analyzed 124,376 White British individuals with no missing data in age, sex, the top 10 genetic PCs, the history of medication, and two measurements of systolic blood pressures at the initial assessment visit. The final phenotypic values for systolic blood pressures are obtained as the mean of the two measurements at initial assessment visit and adjusted for medication record. For individuals with a history of taking medications to control blood pressure, we increase their systolic blood pressure by 15, as suggested by previous genetic association studies on blood pressure in the UK Biobank White British cohort.^49^ We used the GRCh37 deCODE map developed by Bherer et al. (2017) in the analyses of the UK Biobank data.^50^

We first estimated the genome-wide type I error rate when applying our IBD mapping test on the UK Biobank dataset. We selected random subsets of 1000 individuals and simulated 1000 replicates of phenotypes with no genetic associations using the model ***Y*** = 0.05 * *age* + 0.5 * *sex* + ***g*** + ***ε***, where ***g*** and ***ε***are defined in the same way as the studies on the simulated data, and observed age and sex are included as fixed effects. For each replicate, we conducted genome-wide IBD mapping at 0.1 cM intervals and derived the genome-wide significance threshold based on the random subset of 1000 individuals. We calculated the genome-wide type I error rates based on results of 1000 replicates as described previously.

Next, to analyze the genetic association of systolic blood pressure among White British individuals in UK Biobank, we conducted IBD mapping tests adjusted for age, sex, age squared, their interactions, and the first 10 genetic principal components as fixed effects in the model. We used 2 cM as the haplotype length threshold and 0.5 cM as the trimming threshold in multi-individual IBD clustering. We assessed the test results against the genome-wide multiple testing threshold derived from the genotype data of all 124,376 individuals in the analysis.

We also analyzed the data using FiMAP, a recently proposed IBD mapping test that employs the same variance component model but utilizes a score-type statistic for testing the variance component hypothesis with pairwise IBD segments.^27^ To balance power and computational efficiency, FiMAP recommends an IBD threshold of 3 cM because using a shorter threshold (such as 2 cM) would substantially increase the number of IBD segments, resulting in denser local IBD matrices and significantly higher computational demands.^27^ Following the analysis procedure outlined in Chen et al. (2023), we used hap-ibd to identify IBD segments of at least 3 cM in length and performed the FiMAP test at 3,341 consecutive, non-overlapping 1 cM regions across the genome, adjusting the same set of fixed effects as in our IBD mapping test. FiMAP p-values are evaluated against a Bonferroni -adjusted significance threshold of 0.05/^33^41 = 1.5 × 10^−5^.

## Results

### Genome-wide multiple-testing correction

We derived the genome-wide multiple testing threshold for IBD mapping tests on the simulated sequence data and the simulated SNP array data using different haplotype length and trimming thresholds for multi-individual IBD clustering. For each test scenario, the estimated decay parameter 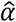 of the modified OU process (see Methods) is shown in Fig S1 along with its 95% bootstrap confidence interval. The 95% genome-wide multiple testing thresholds for test p-values are summarized in Table 1.

**Table 1.** Estimated decay parameters 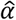 and the corresponding genome-wide 95% significance threshold for test p-values under different haplotype length thresholds (***L***) and trimming thresholds (***T***) for multi-individual IBD detection.

| *L* |  | 2cM |  |  | 3cM |  |  |
| --- | --- | --- | --- | --- | --- | --- | --- |
| *T* |  | 0.25 cM | 0.5 cM | 1 cM | 0.25 cM | 0.5 cM | 1 cM |
| Simulated<br>sequence<br>data | $\hat{\alpha}$ | 1.03 | 1.17 | 1.29 | 0.39 | 0.46 | 0.72 |
| | Threshold | $2.9 \times 10^{-6}$ | $2.8 \times 10^{-6}$ | $2.7 \times 10^{-6}$ | $5.2 \times 10^{-6}$ | $4.7 \times 10^{-6}$ | $3.5 \times 10^{-6}$ |
| Simulated<br>SNP array<br>data | $\hat{\alpha}$ | 1.02 | 1.14 | 1.3 | 0.41 | 0.48 | 0.73 |
| | Threshold | $3.0 \times 10^{-6}$ | $2.8 \times 10^{-6}$ | $2.7 \times 10^{-6}$ | $5.0 \times 10^{-6}$ | $4.5 \times 10^{-6}$ | $3.5 \times 10^{-6}$ |

For each simulated test scenario, the estimated 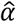 from the simulated sequence data closely matched those from the simulated SNP array data. Using a 3 cM haplotype length threshold for detecting multi-individual IBD resulted in smaller 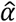 compared to using a 2 cM threshold. At the same haplotype length threshold, increasing the trimming threshold for IBD clustering led to higher 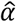 estimates. Larger 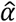 values corresponded to smaller genome-wide multiple-testing thresholds for p-values.

We calculated the theoretical correlations between random variables in each modified OU process using equation (6) and the estimated 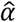 for each test scenario. Across all test scenarios, the observed correlations between test statistics were consistent with the theoretical correlations of the modified OU process for both the simulated sequence data (Fig S2) and the simulated SNP data (Fig S3), especially for pairs of test statistics less than 1 cM apart. These results confirm that the modified OU process provides a reliable approximation to the correlation structure of the IBD mapping test statistics.

For the UK Biobank SNP array data, we estimated the genome-wide multiple testing threshold for IBD mapping tests using a haplotype length threshold of 2 cM and a trimming threshold of 0.5 cM in multi-individual IBD clustering. The estimated decay parameter 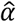 of the corresponding modified OU process was 1.62, with a 95% bootstrap confidence interval of 1.35 and 1.92. The 95% bootstrap confidence interval of the observed correlations overlapped with the trajectory of theoretical correlation of the modified OU process with 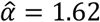 (Fig S4). The estimated multiple-testing p-value threshold for a 5% genome-wide type I error rate was ^2^.^3^ × 10^−6^.

### Genome-wide type I error rates

Table 2 summarizes the estimated genome-wide type I error rates of our IBD mapping test using the proposed genome-wide multiple testing adjustment on both the simulated sequence data and the UK Biobank SNP array data when different haplotype length and trimming thresholds are used for multi-individual IBD detection. As a comparison, we also calculated the proportion of genome-wide IBD mapping replicates where at least one test exceeded the significance threshold based on the Bonferroni correction, which is 0.05/^3^0000 for IBD mapping tests on the simulated sequence data and 0.05/^33^498 for tests on the UK Biobank data.

**Table 2.** Genome-wide type I error rates for the proposed genome-wide multiple-testing adjustment and the Bonferroni adjustment for IBD mapping tests at a genome-wide significance level of 0.05. Results are based on simulated phenotypes under the null hypothesis of no genetic association, using both the simulated sequence data and the UK Biobank SNP-array data, with different haplotype length thresholds (***L***) and trimming thresholds (***T***) for multi-individual IBD detection.

| | $L$ | 2cM | | | 3cM | | |
| --- | --- | --- | --- | --- | --- | --- | --- |
| | $T$ | 0.25 cM | 0.5 cM | 1 cM | 0.25 cM | 0.5 cM | 1 cM |
| Simulated data | Proposed adjustment | 0.046 | 0.057 | 0.065 | 0.049 | 0.064 | 0.059 |
|  | Bonferroni adjustment | 0.03 | 0.04 | 0.046 | 0.019 | 0.026 | 0.032 |
| UK Biobank SNP-array data | Proposed adjustment | 0.04 | 0.035 | 0.052 | 0.065 | 0.048 | 0.038 |
|  | Bonferroni adjustment | 0.028 | 0.023 | 0.033 | 0.024 | 0.026 | 0.018 |

In most test scenarios, our proposed genome-wide multiple-testing adjustment maintains a well-controlled genome-wide type I error rate close to the nominal level of 5%. In contrast, the genome-wide type I error rates based on the Bonferroni threshold are lower than the nominal level, especially with a 3 cM haplotype length threshold, indicating that as expected the Bonferroni correction is conservative.

### Power

The power of our IBD mapping test to detect different types of causal variants, calculated using the corresponding genome-wide significance threshold from Table 2 under various parameter settings for multi-individual IBD clustering, is compared in Figure 1. Our IBD mapping test demonstrates similar power to detect all classes of causal variants when applied to both the simulated sequence data and the simulated SNP array data.

**Figure 1.**
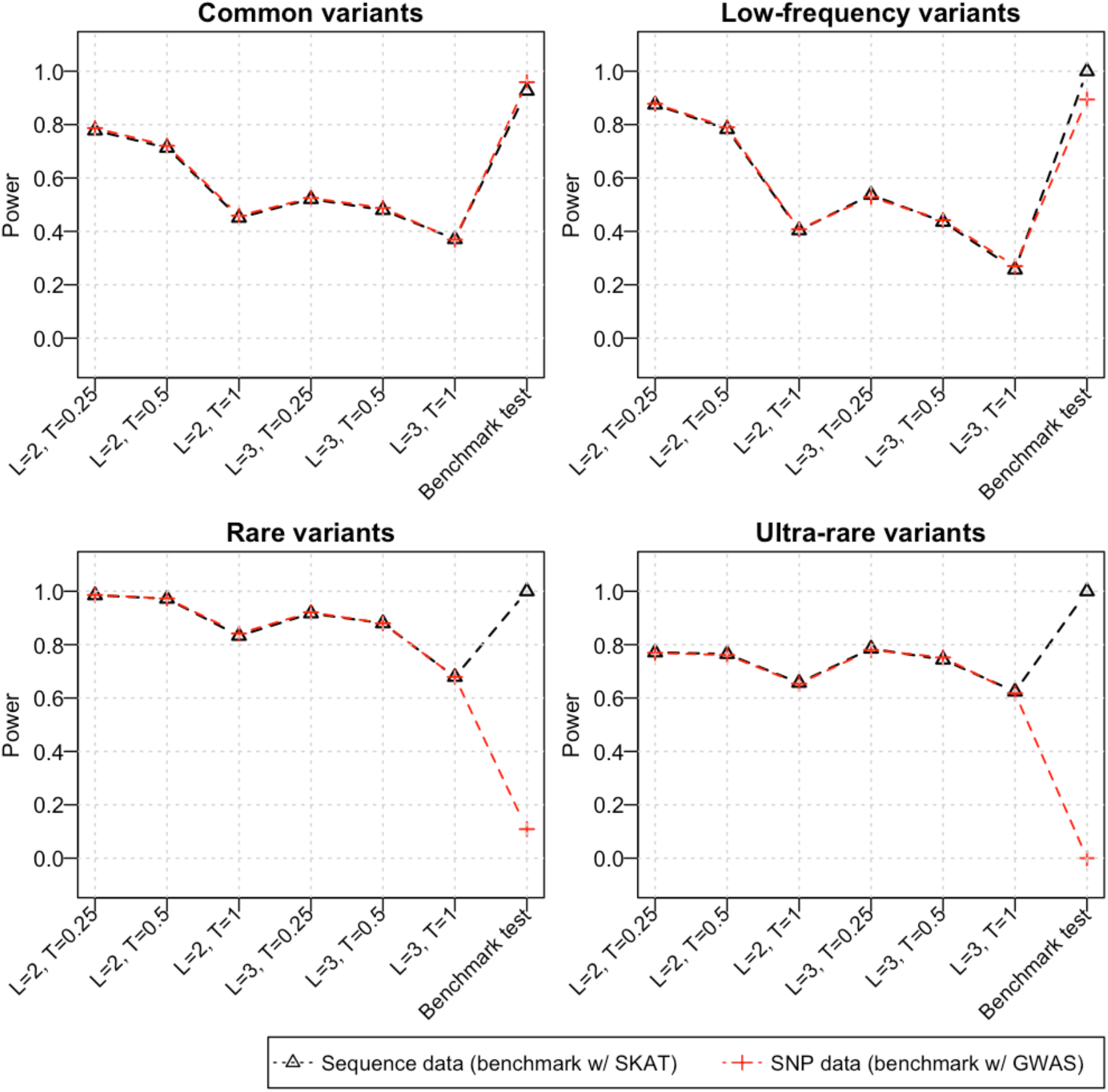
The power of our IBD mapping test for detecting common (>10% MAF), low-frequency (1-10% MAF), rare (0.05-1% MAF), and ultra-rare (<0.05% MAF) causal variants using the proposed genome-wide multiple testing threshold. On the simulated sequence data, we compared the IBD mapping test to the sequence kernel association test (SKAT). On the simulated SNP array data, we compared the IBD mapping test to the traditional single-variant test used in GWAS. The tick labels on the x-axis indicates the haplotype length threshold (***L***) and the trimming threshold (***T***) used for multi-individual IBD detection in the corresponding IBD mapping test.

Across all parameter settings for multi-individual IBD clustering, the highest power to detect causal variants is achieved for rare variants with MAFs between 0.05% and 1%. When using multi-individual IBD detected from haplotypes of length at least 2 cM, trimmed by 0.25 cM or 0.5 cM, the IBD mapping test reaches over 97% power to detect rare variants associated with the simulated phenotypes in both the simulated sequence and SNP-array data. These settings also yield good power (70%-80%) to detect common, low-frequency, or ultra-rare variants. Additionally, the IBD mapping test using a minimum haplotype length of 3 cM with trimming thresholds of 0.25 cM or 0.5 cM for multi-individual IBD clustering also provide good power for detecting rare and ultra-rare variants.

When using the simulated sequence data as the testing dataset, the IBD mapping test does not outperform the SKAT test, which achieved 90% power for detecting common causal variants and 100% power for detecting the other three types of causal variants. When performing tests on the simulated SNP-array data, where none of the causal variants are directly available, the single-variant test achieved higher power for detecting common variants (96%). However, the single-variant test showed similar power to detect low-frequency variants as the IBD mapping test, and it achieved only 11% power for detecting rare variants and completely failed to detect ultra-rare variants. Across all parameter settings, the IBD mapping test significantly outperformed the single-variant test in detecting rare and ultra-rare causal variants. The highest power for detecting rare causal variants reaches 98.6% when using a haplotype threshold of 2 cM and a trimming threshold of 0.25 cM for multi-individual IBD detection, and 78% for detecting ultra-rare causal variants when a haplotype threshold of 3 cM and a trimming threshold of 0.25 cM are used for multi-individual IBD clustering.

Lastly, we evaluated the power gain achieved by the proposed genome-wide multiple-testing adjustment. We calculated the power of our IBD mapping test using the naïve Bonferroni correction 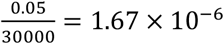 and compared it to results obtained using our proposed genome-wide multiple-testing adjustment on both the simulated sequence data and the simulated SNP array data (Fig S5). In most test scenarios, we observed an absolute increase of at least 2-3% in power using our proposed multiple-testing adjustment compared to using the Bonferroni correction on both the simulated sequence data and the simulated SNP data. Notably, when using a haplotype threshold of 3 cM and a trimming threshold of 0.25 cM for multi-individual IBD clustering, the power to detect low-frequency causal variants is improved by about 8% (from 45.8% to 53.7%) on simulated sequence data and by about 6% (from 47.0% to 52.8%) on simulated SNP array data using our proposed multiple-testing adjustment.

### UK Biobank data analysis

For the genome-wide scan of systolic blood pressure from more than 120k White British individuals in the UK Biobank (Fig 2), the IBD mapping test at 17.759 Mb on chromosome 19 (GRCh37) achieved the genome-wide multiple-testing threshold of ^2^.^3^ × 10^−6^ with p-value=^2^.^2^1 × 10^−7^. In addition, a near-significant signal is observed at 3.790 Mb on chromosome 22 (p-value=^2^.51 × 10^−6^).

**Figure 2.**
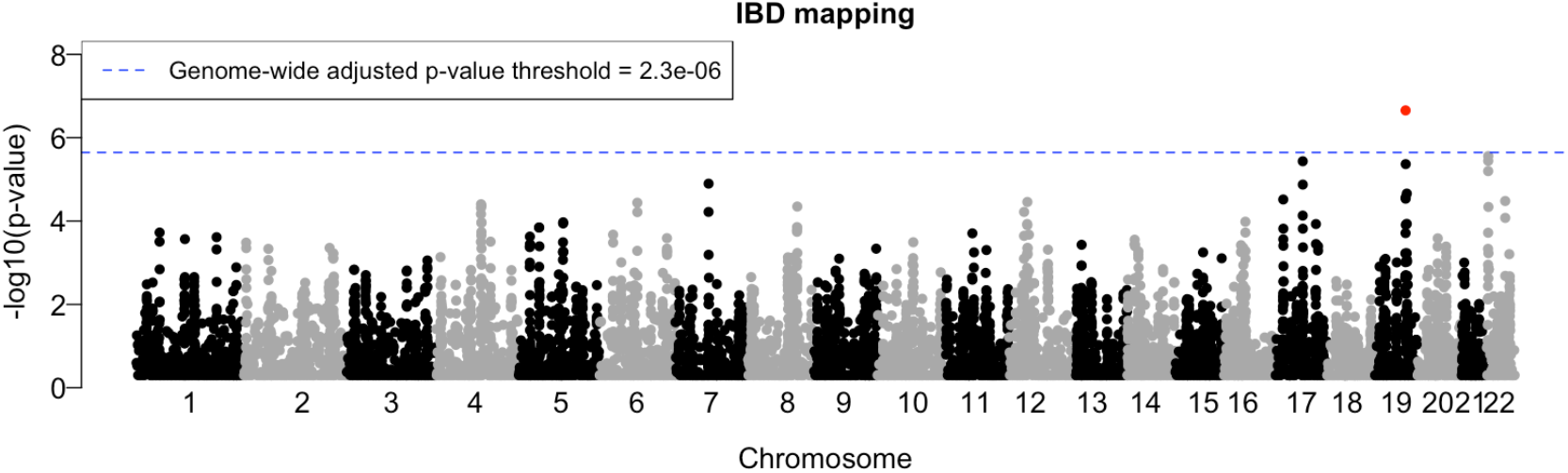
Genome-wide IBD mapping on systolic blood pressure data from 120k White British individuals in the UK Biobank. The negative log10 transformed p-values are plotted against tested positions on each chromosome along the genome. The result that achieved the genome-wide multiple-testing threshold is highlighted in red.

For comparison, we applied the FiMAP test to this dataset at 1 cM intervals across the genome (Fig S6). No significant signals were observed relative to the Bonferroni p-value threshold of 0.05/^33^6^3^ = 1.5 × 10^−5^.

## Discussion

In this study, we developed an IBD mapping approach that leverages multi-individual IBD sharing among distantly related individuals in large, outbred populations. Our approach constructs local relatedness matrices from multi-individual IBD sharing and employs a likelihood ratio statistic to assess the contribution of local genetic similarities to the total phenotypic variation. To address genome-wide multiple-testing, we proposed a p-value adjustment obtained by modeling the correlations between test statistics across the genome.

Through simulation studies, we evaluated the performance of our IBD mapping test across different types of testing datasets and a range of parameter settings for multi-individual IBD clustering. Our results demonstrated that when appropriate haplotype length thresholds and trimming thresholds are chosen for multi-individual IBD detection, the IBD mapping test maintains a well-controlled genome-wide type I error rate and exhibits robust power to detect various types of causal variants, with particular strength in detecting rare or untyped variants compared to traditional GWAS single-variant tests. The power of IBD mapping was similar when applied to both SNP array and sequence data, making it a versatile approach that is suitable for use with either type of dataset. The proposed genome-wide multiple testing adjustment further improved power for our IBD mapping test compared to the Bonferroni threshold.

However, in our simulation study, the IBD mapping test did not outperform the SKAT test when applied to sequence data in which the causal variants were genotyped. It was also less powerful than single-variant GWAS for variants with MAF>10% on SNP array data. These findings emphasize that the strengths of our method lie in its complementary nature rather than as a replacement for existing approaches. Nevertheless, the IBD mapping test may still provide additional perspectives in scenarios involving structural variants or other genetic features that are not directly captured in the sequence data.

Another strength of this study is the genome-wide multiple-testing adjustment we proposed, which serves as an alternative to the commonly used Bonferroni correction. In previous studies, resampling methods such as permutation tests have been employed to derive multiple-testing adjustments that account for the correlations between test statistics^18,51,52^. However, such approaches are computationally intensive, especially in large-scale genome-wide association studies. In contrast, our proposed analytical adjustment is more computationally efficient while still ensuring reliable control of the family-wise type I error rate by addressing the inherent correlation structure in test statistics. This feature is particularly advantageous for large-scale studies where computational efficiency is crucial.

Applications of our IBD mapping approach to the UK Biobank systolic blood pressure data revealed a locus on chromosome 19 with IBD mapping p-value that surpassed the genome-wide threshold. This location is close to the *RCN3* gene on chromosome 19, which has been found to harbor low-frequency and rare variants associated with blood pressure traits.^53,54^ This result illustrates the potential of our method for uncovering biologically relevant signals in large-scale datasets. In comparison, the FiMAP test did not find any significant signals in the same dataset.

However, our method required hundreds of hours to run, whereas FiMAP completed in just a couple of hours, indicating a trade-off between sensitivity and computational cost. While the FiMAP test is well-suited for efficient genome-wide scans in large-scale biobank cohorts, our method may be more advantageous for smaller datasets, where the higher power it offers is more attainable given manageable computational cost.

The performance of our IBD mapping approach is inherently dependent on the quality of the inferred multi-individual IBD sharing. The accuracy and resolution of multi-individual IBD detection, influenced by parameters such as haplotype length and trimming thresholds, directly impact both the type I error rate and the power of the mapping tests. Suboptimal parameter choices or inaccuracies in IBD clustering can lead to loss of power. This suggests a need for further refinement of selection strategies for parameters used for multi-individual IBD detection to balance sensitivity, error control and computational efficiency in our IBD mapping approach.

In addition to the challenges related to multi-individual IBD detection, our study also highlights several areas for further improvement. First, our method requires a significant amount of computational time on biobank-scale cohorts, limiting its scalability to sample size compared to studies using simpler statistical tests, such as single-variant GWAS or SKAT-type tests. These tests generally rely on more straightforward calculations, such as comparing summary statistics, fitting linear models, or computing score-type statistics under the null hypothesis, which are computationally efficient even for large sample sizes. In contrast, our approach is based on a likelihood ratio framework, where optimizing model likelihoods is computationally intensive and becomes increasingly demanding as the sample size grows, thus restricting the scalability of our method.

Furthermore, the resolution of the significant signals identified by our IBD mapping approach was relatively low, because IBD status changes slowly along the chromosome. This suggests that future work could focus on methods such as haplotype testing to refine the IBD signals and pinpoint the causal variants under complex traits.^55,56^ Another limitation of our current approach is its focus on quantitative traits with continuous values; future research extending this method to binary or categorical traits would be useful. Likewise, extending this method to test gene-environment interaction effects by incorporating interaction terms into the model framework could further enhance its applicability for complex trait analyses.

In summary, the IBD mapping approach developed in this study offers a versatile and powerful tool for detecting associations between genomic regions and complex traits, particularly those involving rare or untyped variants. Its robust performance across both SNP array and sequence data underscores its broad applicability to diverse datasets. Future improvements in computational efficiency, integration with higher-resolution methods, and extensions to binary traits, gene-environment interactions, and other complex genetic features could further enhance the utility of identity-by-descent information in advancing our understanding of the genetic architecture underlying complex traits across diverse populations.

## Acknowledgements

Research reported in this publication was supported by the National Human Genome Research Institute of the National Institutes of Health under award number HG007501. The content is solely the responsibility of the authors and does not necessarily represent the official views of the National Institutes of Health. This research has been conducted using the UK Biobank Resource under Application Number 19934.

## Supplementary materials

**Figure S1.**
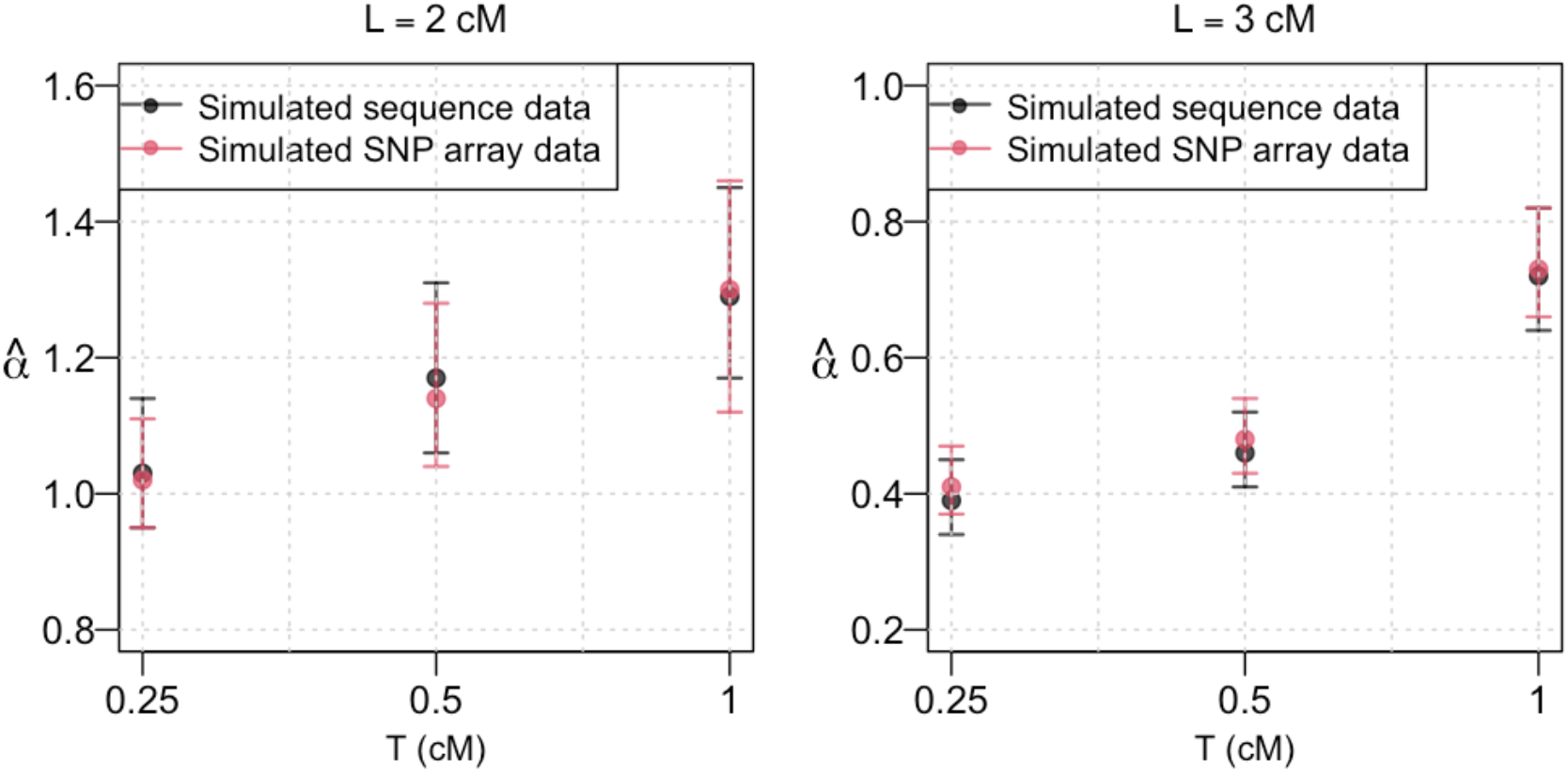
Estimated decay parameter 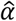 for genome-wide IBD mapping tests under different algorithm parameters for multi-individual IBD detection. The left panel shows results using a haplotype length threshold of 2 cM, while the right panel shows results for a 3 cM threshold. The y-axis represents the estimated 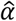, plotted against the corresponding trimming thresholds used in each test. Results from tests on the simulated sequence data are shown in black, and results from tests on the simulated SNP array data are shown in red.

**Fig S2.**
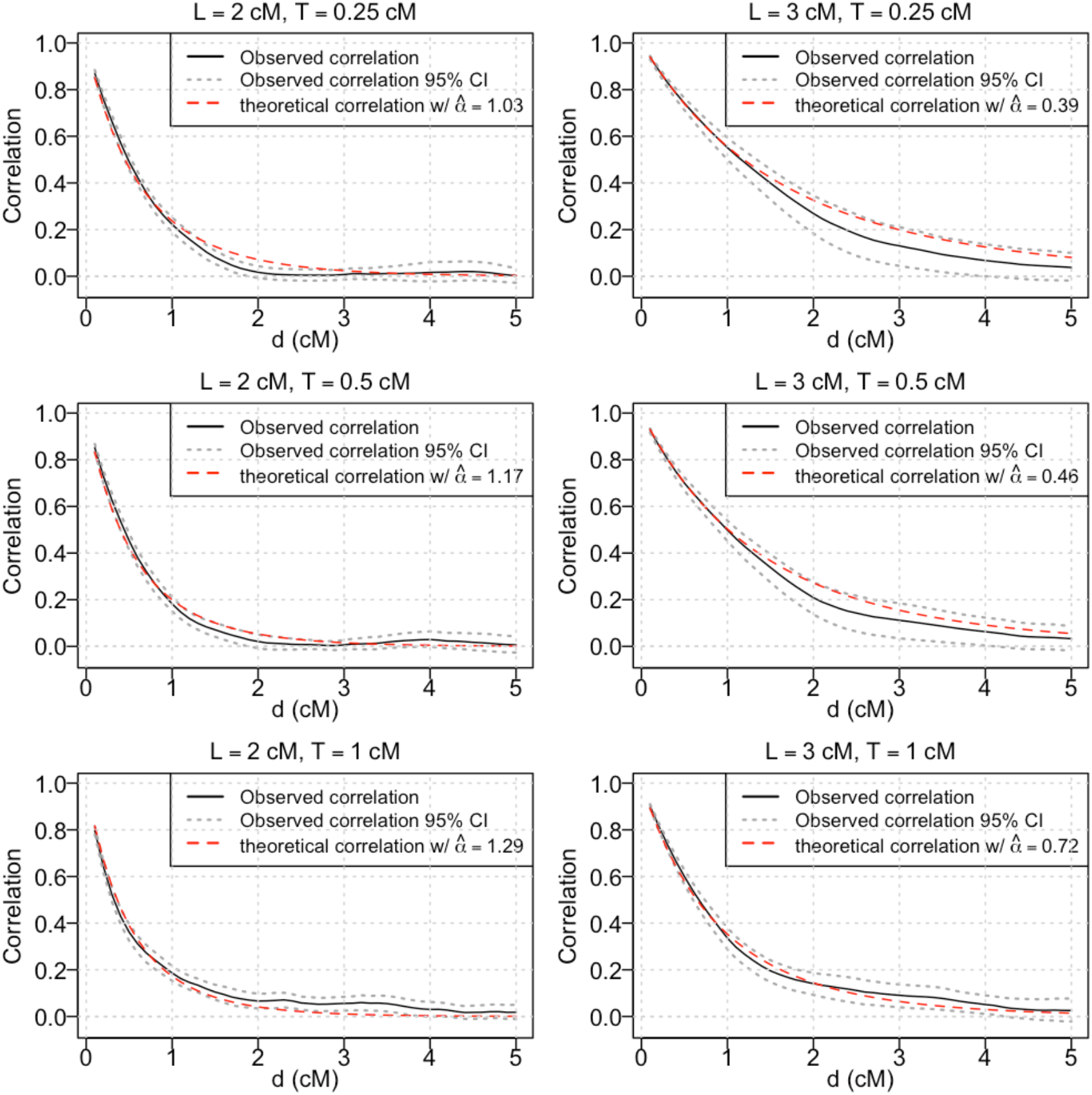
Empirical correlation between test statistics spaced at *d* cM from genome-wide IBD mapping tests with phenotypes simulated under the null hypothesis on simulated sequence data, using different haplotype length thresholds (***L***) and trimming thresholds (***T***) for multi-individual IBD detection. The 95% confidence interval for the correlation between test statistics spaced at *d* cM is constructed by taking the 2.5^th^ and 97.5^th^ percentile of the empirical correlation of test statistics spaced at *d* cM across all bootstrap samples. The theoretical correlation is calculated using the corresponding estimated decay parameter 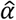.

**Fig S3.**
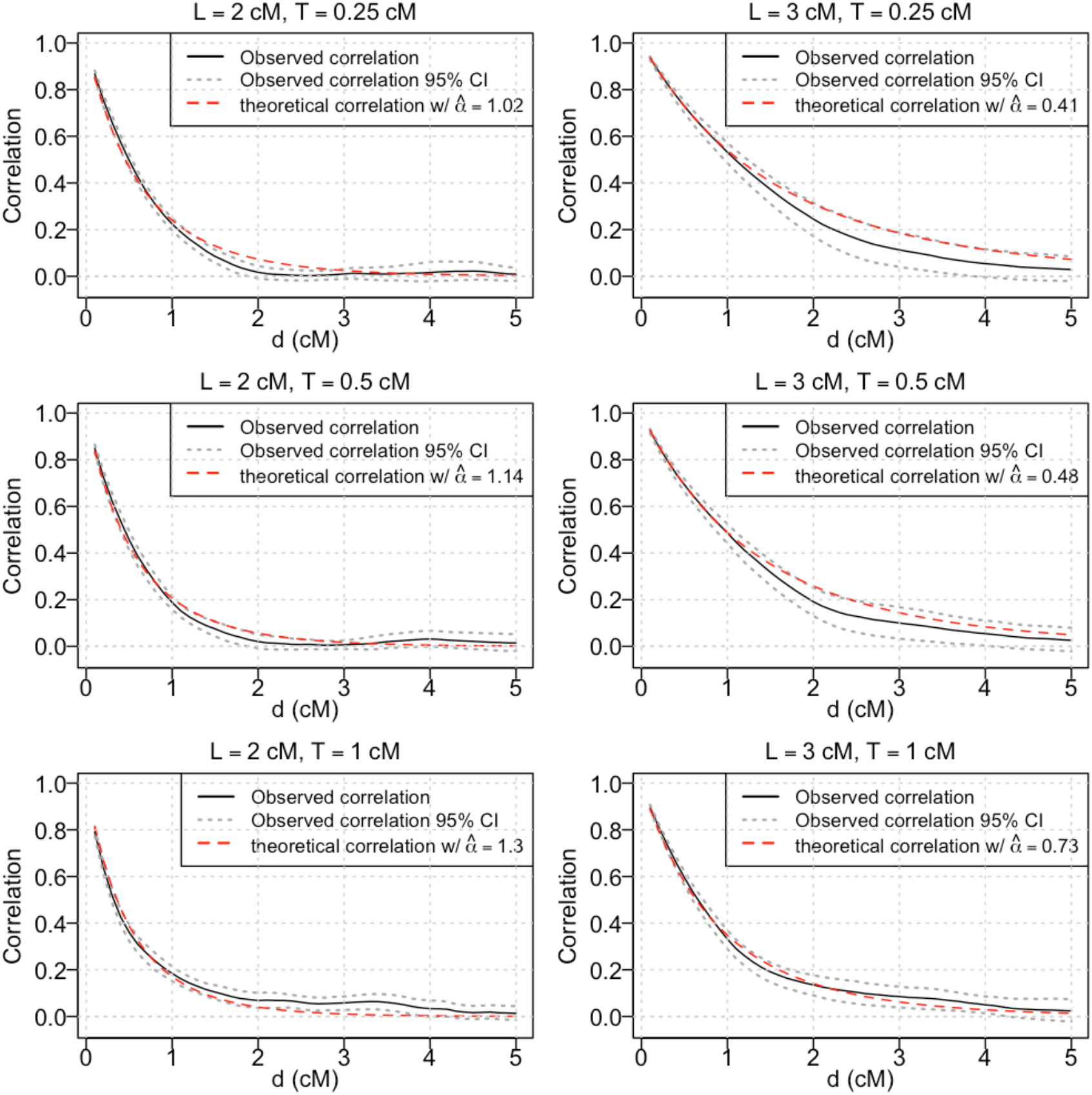
Empirical correlation between test statistics spaced at *d* cM from genome-wide IBD mapping tests with phenotypes simulated under the null hypothesis on simulated SNP array data, using different haplotype length thresholds (***L***) and trimming thresholds (***T***) for multi-individual IBD detection. The 95% confidence interval for the correlation between test statistics spaced at *d* cM is constructed by taking the 2.5^th^ and 97.5^th^ percentile of the empirical correlation of test statistics spaced at *d* cM across all bootstrap samples. The theoretical correlation is calculated using the corresponding estimated decay parameter 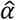.

**Fig S4.**
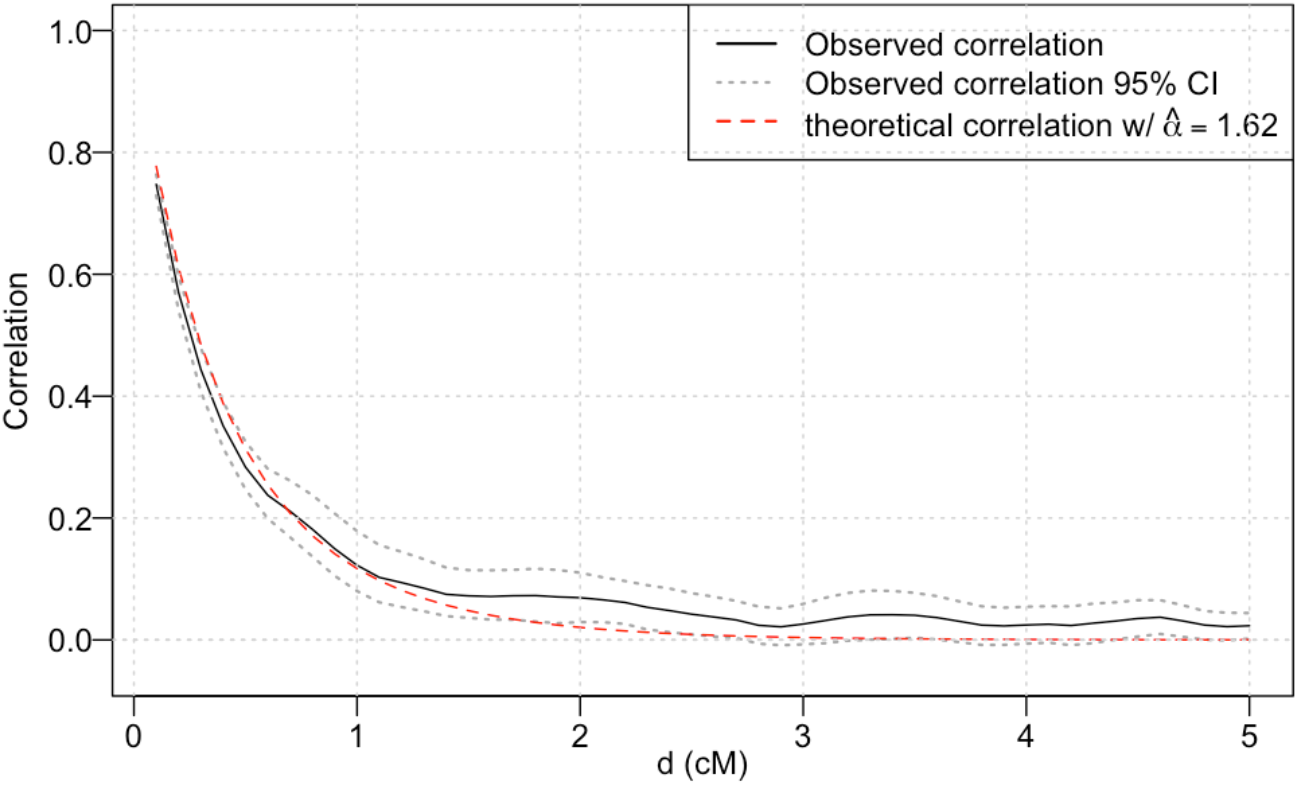
Empirical correlation between test statistics from genome-wide IBD mapping tests with phenotypes simulated under the null hypothesis on the UK Biobank SNP array data, using a haplotype length threshold of 2 cM and a trimming threshold of 0.5 cM. The empirical correlation and their 95% bootstrap confidence intervals are compared to the theoretical correlation calculated with estimated decay parameter 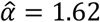.

**Fig S5.**
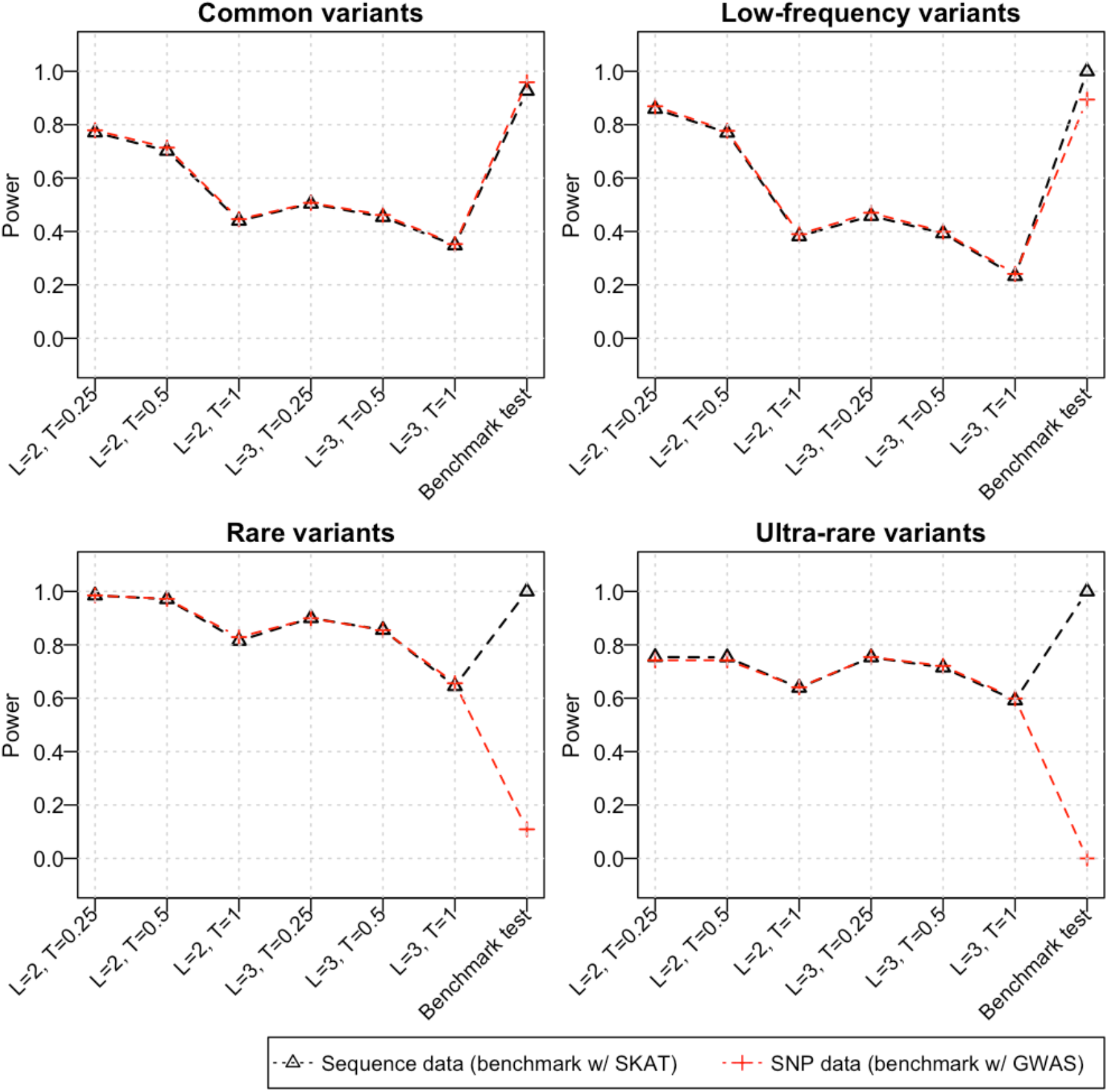
The power of our IBD mapping test for detecting common (>10% MAF), low-frequency (1-10% MAF), rare (0.05-1% MAF), and ultra-rare (<0.05% MAF) causal variants using the Bonferroni correction. On the simulated sequence data, we compared the IBD mapping test to the sequence kernel association test (SKAT). On the simulated SNP array data, we compared the IBD mapping test to the traditional single-variant test used in GWAS. The tick labels on the x-axis indicate the haplotype length threshold (***L***) and the trimming threshold (***T***) used for multi-individual IBD detection in the corresponding IBD mapping test.

**Figure S6.**
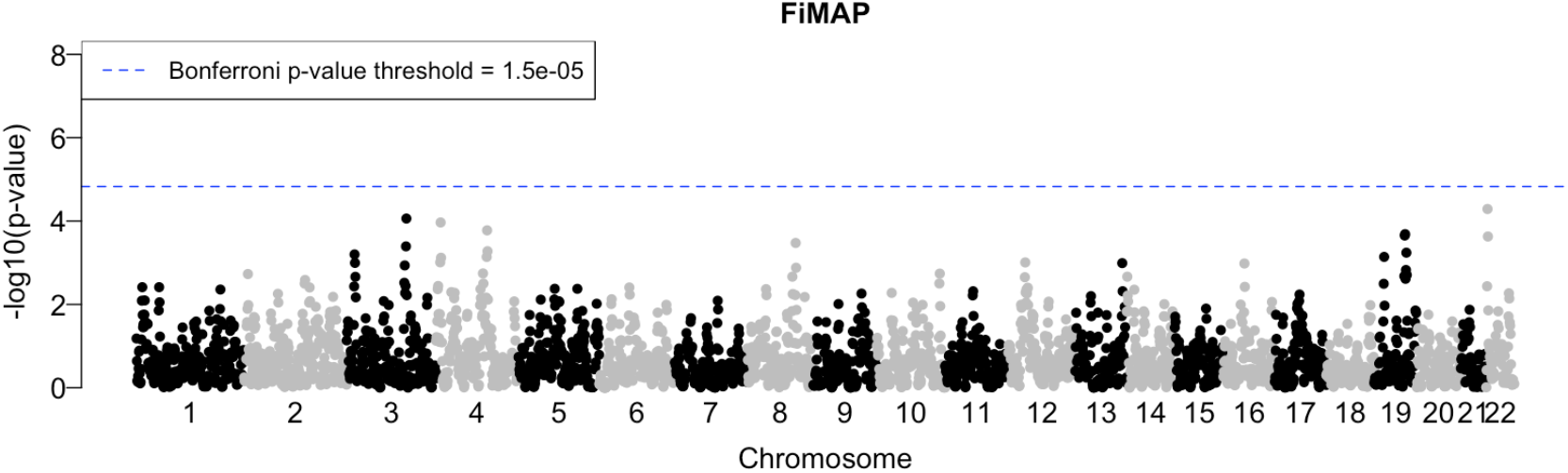
Results of applying the FiMAP test at 1 cM intervals across the genome to systolic blood pressure data from 120k White British individuals in the UK Biobank. Negative log10 transformed p-values are plotted against tested positions on each chromosome along the genome.

## Appendices

## Appendix A. The Null distribution of *W*_*l*_

We consider the mixed-effects model

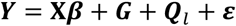

- ***Y*** is a vector of length *N*, where *N* is the sample size
- **X** is a *N* × *p* matrix, where *p* is the number of fixed-effects covariates
- ***β*** is a vector of length *p*
- 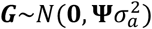, **ψ** is the *N* × *N* genome-wide relatedness matrix
- 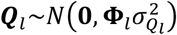, **Φ**_*l*_ is the *N* × *N* local relatedness matrix at a genomic location *l*
- 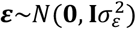 , **I** is the *N* × *N* identity matrix

Denote

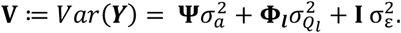

The restricted log likelihood of this model has the following form:

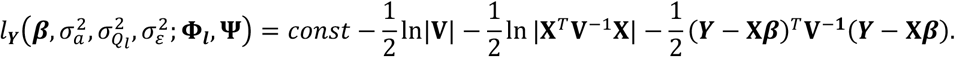

Denote 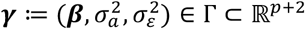 and 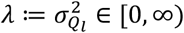. We are interested in testing the hypothesis:

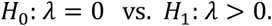

Note that *λ* = 0 is a boundary point of its parameter space.

For each fixed *λ* ≥ 0, define

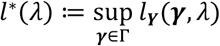

In terms of *l*^*^(*λ*), the test statistic *W*_*l*_ can be written as:

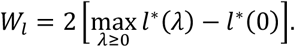

Assume *l*_***Y***_(*γ, λ*) is sufficiently smooth and satisfies usual regularity and identifiability conditions. Denote

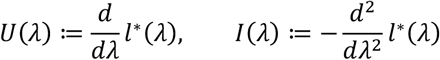

Note that the solution of 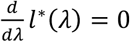 is the unconstrained maximum likelihood estimator, i.e.,

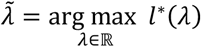

Thus,

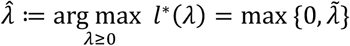

Consider the first derivative of *l*^*^(*λ*) at 0:

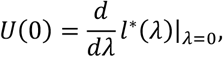

we see

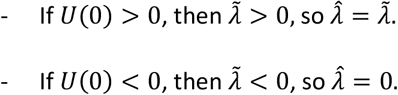

Note that 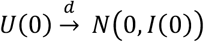. Thus, 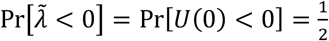.

When 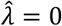, *W*_*l*_ = 0. When 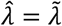, by asymptotic properties of the maximum likelihood estimator, we have that 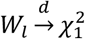. Therefore, the distribution of *W* is a 50:50 mixture of a point mass at 0 and a 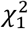 under *H*_0_, as shown in Chernoff (1954), Miller (1977), and Self and Liang (1987).^57-59^

## Appendix B. Correlation structure of the modified OU process

Consider 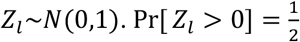, and 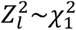. From results shown in Appendix A, it is easily seen that 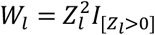.

Now suppose that observations 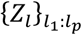 are drawn from an Orenstein-Uhlenbeck process, where

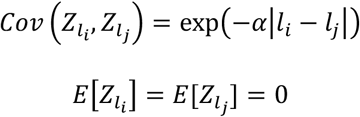

for any two observations 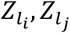 taking at locations *l*_*i*_ and time *l*_*i*_ from this process. We refer to 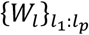 as the modified OU process, i.e., for *l*_*i*_ ∈ {*l*_1_, *l*^2^, … , *l*_*p*_},

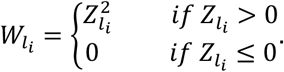

Our goal is to derive 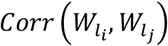.

First, we derive 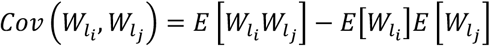. Since the marginal distribution of 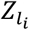 is standard normal, we have:

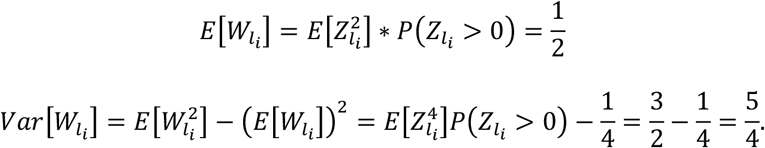

Furthermore,

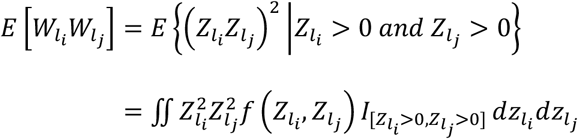

where 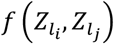 is the probability density function of the joint binormal distribution of 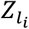 and 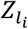. We evaluated this integral using Mathematica^60^, and we obtained that

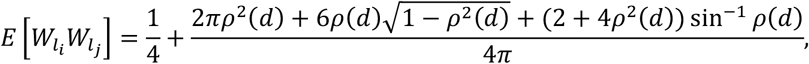

where we define *d* ≔ t*l*_*i*_ − *l*_*j*_t and *ρ*(*d*) ≔ exp(−α*d*). Hence,

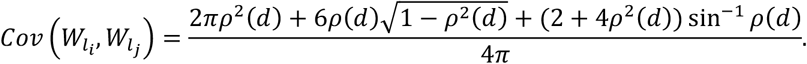

And so

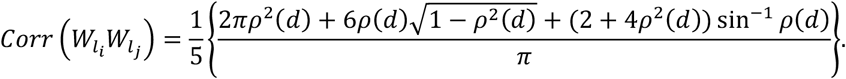

## Appendix C. Simulation algorithm of the modified OU process along a chromosome

### Input

- chrlen: total length of chromosome in centiMorgan
- alpha: value of the decay parameter α
- stepsize: size of the interval between consecutive obervations
- n_iter: total number of replicates generated

### Output

- maxval: a vector of maximum simulated observations in each of the replicates

### Initialization

- Initialize x as an array of n_iter random normal values
- Initialize modified_x as an array where:

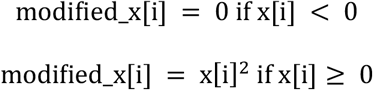
- Set maxval = modified_x

### Iterative process

For *i* from 2 to 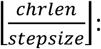

- newx = array of n_iter random normal values with:

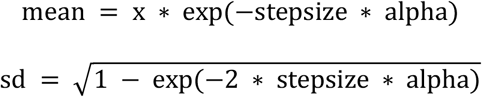
- Initialize new_modified_x as an array where:

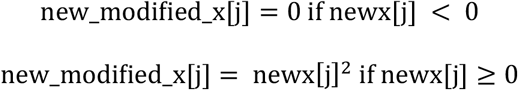
- Update maxval as the element-wise maximum of maxval and new_modified_x
- Set x = newx

Return maxval

## Notes

### Competing Interest Statement

The authors have declared no competing interest.

## References

1. Amos CI. Robust variance-components approach for assessing genetic linkage in pedigrees. American journal of human genetics. 1994;54(3):535.

2. Blangero J, Almasy L. Multipoint oligogenic linkage analysis of quantitative traits. Genetic epidemiology. 1997;14(6):959–964.

3. Almasy L, Blangero J. Multipoint quantitative-trait linkage analysis in general pedigrees. The American Journal of Human Genetics. 1998;62(5):1198–1211.

4. Blangero J, Williams JT, Almasy L. Quantitative trait locus mapping using human pedigrees. Human Biology. 2000:35–62.

5. Day-Williams AG, Blangero J, Dyer TD, Lange K, Sobel EM. Linkage analysis without defined pedigrees. Genetic epidemiology. 2011;35(5):360–370.

6. Hirschhorn JN, Daly MJ. Genome-wide association studies for common diseases and complex traits. Nature reviews genetics. 2005;6(2):95–108.

7. McCarthy MI, Hirschhorn JN. Genome-wide association studies: potential next steps on a genetic journey. Human molecular genetics. 2008;17(R2):R156–R165.

8. Visscher PM, Wray NR, Zhang Q, et al. 10 years of GWAS discovery: biology, function, and translation. The American Journal of Human Genetics. 2017;101(1):5–22.

9. McCarthy MI, Abecasis GR, Cardon LR, et al. Genome-wide association studies for complex traits: consensus, uncertainty and challenges. Nature reviews genetics. 2008;9(5):356–369.

10. Eichler EE, Flint J, Gibson G, et al. Missing heritability and strategies for finding the underlying causes of complex disease. Nature reviews genetics. 2010;11(6):446–450.

11. Tam V, Patel N, Turcotte M, Bossé Y, Paré G, Meyre D. Benefits and limitations of genome-wide association studies. Nature Reviews Genetics. 2019;20(8):467–484.

12. Madsen BE, Browning SR. A groupwise association test for rare mutations using a weighted sum statistic. PLoS genetics. 2009;5(2):e1000384.

13. Wu MC, Lee S, Cai T, Li Y, Boehnke M, Lin X. Rare-variant association testing for sequencing data with the sequence kernel association test. The American Journal of Human Genetics. 2011;89(1):82–93.

14. Lee S, Emond MJ, Bamshad MJ, et al. Optimal unified approach for rare-variant association testing with application to small-sample case-control whole-exome sequencing studies. The American Journal of Human Genetics. 2012;91(2):224–237.

15. Liu Y, Chen S, Li Z, Morrison AC, Boerwinkle E, Lin X. ACAT: a fast and powerful p value combination method for rare-variant analysis in sequencing studies. The American Journal of Human Genetics. 2019;104(3):410–421.

16. Purcell S, Neale B, Todd-Brown K, et al. PLINK: a tool set for whole-genome association and population-based linkage analyses. The American journal of human genetics. 2007;81(3):559–575.

17. Gusev A, Kenny EE, Lowe JK, et al. DASH: a method for identical-by-descent haplotype mapping uncovers association with recent variation. The American Journal of Human Genetics. 2011;88(6):706–717.

18. Browning SR, Thompson EA. Detecting rare variant associations by identity-by-descent mapping in case-control studies. Genetics. 2012;190(4):1521–1531.

19. Browning SR, Browning BL. Identity by descent between distant relatives: detection and applications. Annual review of genetics. 2012;46:617–633.

20. Qian Y, Browning BL, Browning SR. Efficient clustering of identity-by-descent between multiple individuals. Bioinformatics. 2014;30(7):915–922.

21. Vacic V, Ozelius LJ, Clark LN, et al. Genome-wide mapping of IBD segments in an Ashkenazi PD cohort identifies associated haplotypes. Human molecular genetics. 2014;23(17):4693–4702.

22. Westerlind H, Imrell K, Ramanujam R, et al. Identity-by-descent mapping in a Scandinavian multiple sclerosis cohort. European Journal of Human Genetics. 2015;23(5):688–692.

23. Belbin GM, Odgis J, Sorokin EP, et al. Genetic identification of a common collagen disease in puerto ricans via identity-by-descent mapping in a health system. Elife. 2017;6:e25060.

24. Hsueh W-C, Nair AK, Kobes S, et al. Identity-by-descent mapping identifies major locus for serum triglycerides in Amerindians largely explained by an APOC3 founder mutation. Circulation: Cardiovascular Genetics. 2017;10(6):e001809.

25. Henden L, Twine NA, Szul P, et al. Identity by descent analysis identifies founder events and links SOD1 familial and sporadic ALS cases. NPJ genomic medicine. 2020;5(1):32.

26. Chen H, Naseri A, Zhi D. FiMAP: A fast identity-by-descent mapping test for Biobank-scale cohorts. Plos Genetics. 2023;19(12):e1011057.

27. Chen H, Naseri A, Zhi D. FiMAP: A fast identity-by-descent mapping test for Biobank-scale cohorts. medRxiv. 2021:2021.06.30.21259773.

28. Browning SR, Browning BL. Biobank-scale inference of multi-individual identity by descent and gene conversion. The American Journal of Human Genetics. 2024;111(4):691–700.

29. He D. IBD-Groupon: an efficient method for detecting group-wise identity-by-descent regions simultaneously in multiple individuals based on pairwise IBD relationships. Bioinformatics. 2013;29(13):i162–i170.

30. Almasy L, Blangero J. Variance component methods for analysis of complex phenotypes. Cold Spring Harbor Protocols. 2010;2010(5):pdb.top77.

31. Page GP, Amos CI, Boerwinkle E. The quantitative LOD score: test statistic and sample size for exclusion and linkage of quantitative traits in human sibships. The American Journal of Human Genetics. 1998;62(4):962–968.

32. Broyden CG. The convergence of a class of double-rank minimization algorithms: 2. The new algorithm. IMA journal of applied mathematics. 1970;6(3):222–231.

33. Fletcher R. A new approach to variable metric algorithms. The computer journal. 1970;13(3):317–322.

34. Goldfarb D. A family of variable metric updates derived by variational means, v. 24. Mathematics of Computation. 1970:21–55.

35. Shanno DF. Conditioning of quasi-Newton methods for function minimization. Mathematics of computation. 1970;24(111):647–656.

36. Byrd RH, Lu P, Nocedal J, Zhu C. A limited memory algorithm for bound constrained optimization. SIAM Journal on scientific computing. 1995;16(5):1190–1208.

37. Virtanen P, Gommers R, Oliphant TE, et al. SciPy 1.0: fundamental algorithms for scientific computing in Python. Nature methods. 2020;17(3):261–272.

38. Shor T, Kalka I, Geiger D, Erlich Y, Weissbrod O. Estimating variance components in population scale family trees. PLoS Genetics. 2019;15(5):e1008124.

39. Davis TA. scikit-sparse: Sparse matrix tools extending scipy.sparse. 2021.

40. Zhou Y, Browning SR, Browning BL. A fast and simple method for detecting identity-by-descent segments in large-scale data. The American Journal of Human Genetics. 2020;106(4):426–437.

41. Zhou Y, Browning SR, Browning BL. IBDkin: fast estimation of kinship coefficients from identity by descent segments. Bioinformatics. 2020;36(16):4519–4520.

42. Lander ES, Botstein D. Mapping mendelian factors underlying quantitative traits using RFLP linkage maps. Genetics. 1989;121(1):185–199.

43. Siegmund D, Yakir B. The statistics of gene mapping. vol 1. Springer; 2007.

44. Grinde KE, Brown LA, Reiner AP, Thornton TA, Browning SR. Genome-wide significance thresholds for admixture mapping studies. The American Journal of Human Genetics. 2019;104(3):454–465.

45. Kelleher J, Lohse K. Coalescent simulation with msprime. Statistical Population Genomics. 2020;986:191–230.

46. Baumdicker F, Bisschop G, Goldstein D, et al. Efficient ancestry and mutation simulation with msprime 1.0. Genetics. 2022;220(3):iyab229.

47. Browning BL, Browning SR. Detecting identity by descent and estimating genotype error rates in sequence data. The American Journal of Human Genetics. 2013;93(5):840–851.

48. Bycroft C, Freeman C, Petkova D, et al. The UK Biobank resource with deep phenotyping and genomic data. Nature. 2018;562(7726):203–209.

49. Warren HR, Evangelou E, Cabrera CP, et al. Genome-wide association analysis identifies novel blood pressure loci and offers biological insights into cardiovascular risk. Nature genetics. 2017;49(3):403–415.

50. Bhérer C, Campbell CL, Auton A. Refined genetic maps reveal sexual dimorphism in human meiotic recombination at multiple scales. Nature communications. 2017;8(1):1–9.

51. Browning BL. PRESTO: rapid calculation of order statistic distributions and multiple-testing adjusted P-values via permutation for one and two-stage genetic association studies. BMC bioinformatics. 2008;9:1–5.

52. Marees AT, De Kluiver H, Stringer S, et al. A tutorial on conducting genome-wide association studies: Quality control and statistical analysis. International journal of methods in psychiatric research. 2018;27(2):e1608.

53. Surendran P, Feofanova EV, Lahrouchi N, et al. Discovery of rare variants associated with blood pressure regulation through meta-analysis of 1.3 million individuals. Nature genetics. 2020;52(12):1314–1332.

54. He KY, Kelly TN, Wang H, et al. Rare coding variants in RCN3 are associated with blood pressure. BMC genomics. 2022;23(1):148.

55. Browning SR. Multilocus association mapping using variable-length Markov chains. The American Journal of Human Genetics. 2006;78(6):903–913.

56. Browning BL, Browning SR. Efficient multilocus association testing for whole genome association studies using localized haplotype clustering. Genetic Epidemiology: The Official Publication of the International Genetic Epidemiology Society. 2007;31(5):365–375.

57. Chernoff H. On the distribution of the likelihood ratio. The Annals of Mathematical Statistics. 1954:573–578.

58. Miller JJ. Asymptotic properties of maximum likelihood estimates in the mixed model of the analysis of variance. The Annals of Statistics. 1977:746–762.

59. Self SG, Liang K-Y. Asymptotic properties of maximum likelihood estimators and likelihood ratio tests under nonstandard conditions. Journal of the American Statistical Association. 1987;82(398):605–610.

60. Wolfram Research I. Mathematica. Version 14.1 ed. Champaign, Illinois: Wolfram Research, Inc.; 2024.

